# Network analysis of the human structural connectome including the brainstem: a new perspective on consciousness

**DOI:** 10.1101/2022.07.26.501537

**Authors:** Salma Salhi, Youssef Kora, Gisu Ham, Hadi Zadeh Haghighi, Christoph Simon

**Affiliations:** Department of Physics and Astronomy, University of Calgary, Calgary, AB T2N 1N4, Canada; Hotchkiss Brain Institute, University of Calgary, Calgary, Canada; Fishtank Consulting, 520 5 Ave SW Suite 2200, Calgary, AB T2P 3R7

## Abstract

The underlying anatomical structure is fundamental to the study of brain networks and is likely to play a key role in the generation of conscious experience. We conduct a computational and graph-theoretical study of the human structural connectome incorporating a variety of subcortical structures including the brainstem, which is typically not considered in similar studies. Our computational scheme involves the use of Python DIPY and Nibabel libraries to develop an averaged structural connectome comprised of 100 healthy adult subjects. We then compute degree, eigenvector, and betweenness centralities to identify several highly connected structures and find that the brainstem ranks highest across all examined metrics. Our results highlight the importance of including the brainstem in structural network analyses. We suggest that structural network-based methods can inform theories of consciousness, such as global workspace theory (GWT), integrated information theory (IIT), and the thalamocortical loop theory.

## Introduction

Network-based approaches are exceedingly useful tools for investigating the computational and informational power of the brain [1]. The core idea involves the reduction of an object as complex and generally intractable as the brain to a much simpler system of basic interacting neuronal elements [2]. This profound simplification has proven to be quite powerful, leading to insights into a variety of neuroscientific topics and applications [3]. For instance, diseases such as Alzheimer’s, schizophrenia, and Parkinson’s disease can be seen as disorders of brain networks [4]. The emerging field of network neuroscience [5] is working towards a theoretical framework which unifies the principles underlying the wide range of observed neurobiological phenomena, and deepens our understanding of the relationship between brain structure and function [6].

The goals of such network-based methods include the mapping, modeling and analysis of brain networks, both at a structural and functional level. Structural brain networks are constructed to reflect the direct fibre connections between the different anatomical regions, while functional networks refer to relationships between components which are statistically correlated regardless of the strength of their anatomical connections [7]. Studies of the latter are quite ubiquitous in the literature. These networks are typically extracted through techniques such as electroencephalography (EEG) and functional magnetic resonance imaging (fMRI), which enable the investigation of the topological properties of the dynamical patterns that emerge, both in the resting state and during the performance of tasks [8, 9, 10]. The study of structural networks, on the other hand, has gained prominence in more recent years, largely owing to the recent advances in non-invasive techniques that allow the imaging and mapping of the structural connectome [11, 12].

The human structural connectome provides a foundation for neurobiological research [13]. Indeed, the structural wiring of the brain constrains the cost of controlling a particular brain state; the transition to higher information content states is more costly and is modulated by the anatomical structure [14]. Functional networks are fundamentally influenced by their structural counterparts; it has been shown that the strength and persistence of functional connectivity is constrained by the anatomical cortex [15]. In fact, it is thought that anatomical connectivity allows for the reconciliation of the opposing requirements for functional networks, integration and segregation [16], where integration refers to the ability to quickly combine information from distant brain regions, and segregation refers to the ability for specialized processing to occur within dense regions [17].

On the topic of consciousness, the anatomies of the thalamus and of the cortex are believed to be essential to the underlying neural mechanisms [18]. When the anatomy of the brain is altered, some disorders of consciousness, such as a vegetative state (VM) or a minimally conscious state (MCS) are sometimes observed. Indeed, it has been shown that these patients tend to exhibit significantly reduced structural connections between the basal ganglia, thalamus, and frontal cortex [19]. Another study found a significant decrease in volume in the globus pallidus and the thalamus, especially prominent in the thalamus [20]. These findings suggest that the specific structural wiring in the brain is important for conscious processing, revealing an important application for the structural connectome.

There are a number of theories of consciousness that have been proposed over the past two decades. Among the most prominent is global workspace theory (GWT), which proposes a “global access hypothesis”, in which a fleeting memory capacity enables back and forth access between separate brain functions [21]. In essence, “global access” to sensory content may be a requirement for consciousness [22], where conscious sensory content is accessed when needed and is distributed to a wide array of networks that are not presently conscious. It is thought that corticocortical and corticothalamic fibers are used to distribute the sensory content that is conscious at a given moment [21]. The global workspace would thus consist of a distributed set of heavily interconnected cortical neurons that send and receive information across long-range axons [23].

Integrated information theory (IIT) is an alternative framework which characterizes consciousness through maximally irreducible conceptual structures (MICS) that specify both the quality of an experience as well as the amount of integrated information associated with it [24]. While IIT acknowledges that anatomical connectivity can influence the MICS [24], the theory is more often examined from a functional connectivtity perspective. However, some evidence currently suggests that the neural correlates of consciousness (NCCs) as identified by IIT are likely located in certain parts of the cortico-thalamic system [25], and the entire cerebral cortex is thought to be involved in the subjective experience, albeit passively.

Another example is thalamocortical loop theory, which relies mostly on functional connectivity analysis, specifically by measuring gamma band activity. It conjectures that the thalamus acts as a hub which modulates communication between different cortical regions [26]. The thalamic dynamic core theory is similar, positing that consciousness arises from synchronized activity in thalamic nuclei, which is driven by cortico-thalamic connections that transfer information [27]. The anatomy of the thalamocortical system is thought to be well suited for the neural mechanisms underlying consciousness [18].

Alternatively, it has been proposed that the brainstem is necessary for “core consciousness” – the simplest form of consciousness and the self – because the somato-sensory nuclei in the brainstem are best suited for its modulation [28]. Some theorists maintain that the regulation and arousal of consciousness are affected by the same part of the brain, which is thought to be the brainstem [29], and some even suggest that the brainstem alone can keep a subject conscious [30]. This provided further motivation for the consideration of the brainstem in our structural network analysis.

Such theories are typically evaluated from a functional perspective, despite making hypotheses about the locations of the physical substrates of consciousness. This motivated us to investigate them from the point of view of structural networks. The goal of this study is to shed new light on the importance of certain anatomical regions within the overall structural network, and provide new insight as to their role within each of these theories of consciousness.

MRTrix is perhaps the most widely used software in brain network studies, but the atlas most readily available to its users is the FreeSurfer Desikan-Killiany atlas, which consists of only 84 structures and does not include the brainstem. The full list of these structures can be found in **S2 Table**. This atlas has been used in studies of structural-functional relationships (see, for instance, [31, 32, 33]). The brainstem is conspicuously missing in this atlas, but there is convincing evidence to suggest that it is important for consciousness. Several studies have shown that brainstem resting-state connectivity decreases under propofol-induced mild sedation (see [34], [35]). There is also evidence to suggest that the cortex may not be necessary for consciousness as long as the brainstem is still intact [30], as mentioned previously. With this in mind, we set out to investigate how the inclusion of the brainstem (and other subcortical structures that are excluded in the Desikan-Killiany atlas) affects the resulting structural connectome. To that end, we designed a computational fibre tractography method in Python using DIPY and Nibabel libraries. Using these algorithms, we were able to extract 104 structures from dMRI images provided by the Human Connectome Project (HCP). We constructed two connectomes: one containing the 104 structures extracted using our Python method, which includes the brainstem, and the other comprising the 84 structures typically found in the default MRTrix Desikan-Killiany atlas. We shall henceforth refer to the former as the “extended connectome”, and the latter as the “restricted connectome”.

With the networks obtained, we identified the most structurally important brain regions as those corresponding to nodes occupying the most central positions in each network, i.e., hubs. There are a number of graph-theoretical measures to characterize the centrality of a given node [36]. We started by the simplest of all, degree centrality (also known as node degree, node weight, or connectivity strength), which is defined as the sum of all connections to the node in question. In the case of a weighted network such as ours, this would simply correspond to the total number of streamline connections associated with each brain region. Such a quantity measures the extent to which a given node has strong connections to other nodes in the network, irrespective of their importance.

The importance of other nodes is taken into account by computing the eigenvector centrality [37, 38], which has been applied in more recent times to fMRI data in the human brain [39, 40]. This measure privileges nodes that are well-connected to other nodes in the network which are themselves strongly connected. As the name suggests, this is accomplished by the eigenvector decomposition of the connectivity matrix.

The third measure we computed for our networks is betweenness centrality [41]. Unlike the aforementioned types of centrality, which directly quantify the connectivity of a given node to the rest of the network, betweenness centrality measures the extent to which the node is strategically located in the network. This is achieved by considering the number of shortest geodesic paths between all possible pairs of nodes that pass through the node of interest. Its computation is of complexity *O*(*n*^3^), which renders it an unfeasible measure for sufficiently large networks. It has, however, been used in the context of brain networks with a moderate number of nodes [42, 43], and since the sizes of the networks considered here do not exceed 104 regions, we did not contend with any serious computational limitations.

## Materials and methods

### Data Analysis

No new data was collected for this project. Data was obtained from the Human Connectome Project (HCP) database. The HCP project (Principal Investigators: Bruce Rosen, M.D., Ph.D., Martinos Center at Massachusetts General Hospital; Arthur W. Toga, Ph.D., University of Southern California, Van J. Weeden, MD, Martinos Center at Massachusetts General Hospital) is supported by the National Institute of Dental and Craniofacial Research (NIDCR), the National Institute of Mental Health (NIMH) and the National Institute of Neurological Disorders and Stroke (NINDS). HCP is the result of efforts of co-investigators from the University of Southern California, Martinos Center for Biomedical Imaging at Massachusetts General Hospital (MGH), Washington University, and the University of Minnesota. We used data from 100 adult subjects ranging in age from 22 to 35 years, with 44 females and 56 males, all from the “WU-Minn HCP Data - 1200 Subjects” round of data collection. Both pre-processed T1-weighted structural images and 3T dMRI images were used in our computational fibre tracking method. Python DIPY and NiBabel libraries were utilized to perform the streamline calculations using a constrained spherical deconvolution model and probabilistic fibre tracking functions, which are built in the libraries.

A white matter mask was first created by extracting labels of structures provided in the HCP structural data. Regions that matched the labels for white matter were identified, and the white matter mask was then used to calculate the seeds from which to begin the fibre tracking. For the extended connectome, the mask included the brainstem and a few other structures commonly omitted, the full list of which is found in **S1 Table**. A response function, which is used to form the constrained spherical deconvolution (CSD) model, was then calculated using the DIPY `auto_response_ssst` function. The CSD model was then combined with the structural white matter mask to generate a generalized fractional anistropy (GFA) model, which would aid in the fibre tracking process. A probabilistic direction getter model was then created using a maximum angle of 45°, and this was finally used along with seeds and the diffusion affine matrix to generate a list of streamlines, using the DIPY `LocalTracking` class. A list of endpoints was generated by extracting the first and last elements in each streamline list. This was then used in conjunction with the label mask to assess which structures each endpoint corresponded to in order to create a bi-directional label map. This is essentially a connectivity matrix that matches the total number of streamlines associated with a structure to the correct endpoint.

Due to the probabilistic nature of this tracking algorithm, the program was run three times for each subject and the results were averaged across all three runs in order to corroborate the results of the runs and ensure accuracy. These averages were then collected and averaged across 100 subjects to produce one overarching averaged connectome, representative of all subjects. All python code files are provided on the Github repository created for this project.

The same method was used to calculate the connectivity matrix with a reduced number of structures. The white matter masks were altered to eliminate undesired structures, enabling the fibre tracking algorithm to only examine the 84 structures in the restricted connectome, the full list of which can be found in **S2 Table** in the appendix. A list of the 20 eliminated structures can be found in **S3 Table**.

### Centrality Measures

We compute degree centrality simply as the sum of all connections to a given node

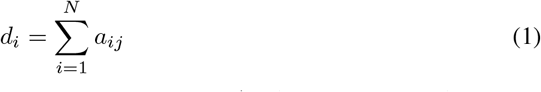

where *a*_*ij*_ are the elements of the structural connectivity matrix *A*. The next examined metric is eigenvector centrality, defined as the components of the eigenvector **v** corresponding to the largest eigenvalue of *A*. The signficance of such a quantity may be seen by looking at the eigenvalue equation

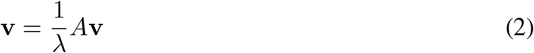

where *λ* is the largest eigenvalue of *A*. By expanding the matrix multiplication process, one may write each component of **v** as

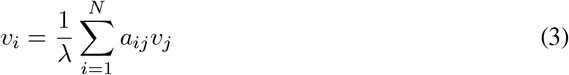

which is similar to Eq 1, but the sum is now weighted by the scores of the node. These scores are all guaranteed to be positive by the Perron-Frobenius theorem, provided that *A* is non-negative and irreducible, i.e., it has at least one non-zero off-diagonal element in each row and column (which will always be true in the cases of interest). These values are only unique up to an overall multiplication factor which depends on the choice of normalization [44], but such a factor will not play a role in the process of comparing nodes within the same network, which is our goal in this study. Finally, betweenness centrality is defined as

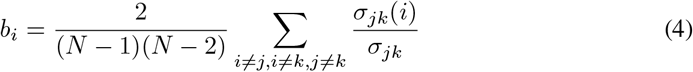

where *σ*_*jk*_ is the number of shortest paths between nodes *j* and *k*, and *σ*_*jk*_(*i*) is the number of shortest paths between nodes *j* and *k* that pass through node *i*. To define the shortest paths in a complete weighted graph such as ours, a notion of distance is required, which was simply taken to be the reciprocal of the weight.

Together, *d*_*i*_, *v*_*i*_, and *b*_*i*_ constitute our node centrality metrics. By studying all three, we compare the graph-theoretical importance of the various brain regions that comprise the connectome, and thus are able to estimate their relative influence within the structural connectome.

## Results

We constructed two connectomes, an extended one containing 104 structures, and a restricted one following the Desikan-Killiany atlas, which eliminates 20 structures from the extended connectome. The connectomes are depicted in Fig 1 and Fig 2 respectively. With the absence of the brainstem in the restricted connectome, the connectome appears less well-connected, with the central hub seen in Fig 1 clearly much less prominent in Fig 2. We can more clearly see this distinction when comparing the connectivity plots. When the 20 subcortical structures are eliminated in the restricted connectome (see table **S3 Table** for a complete list), many of the strong subcortical-subcortical connections (upper left portion of Fig 3a) disappear. The large white squares in Fig 3b reflect a large reduction in the number of subcortical-cortical connections, which has substantially altered the connectivity matrix. The full lists of the extended and restricted structures can be found in **S1 Table** and **S2 Table** respectively.

**Fig 1.**
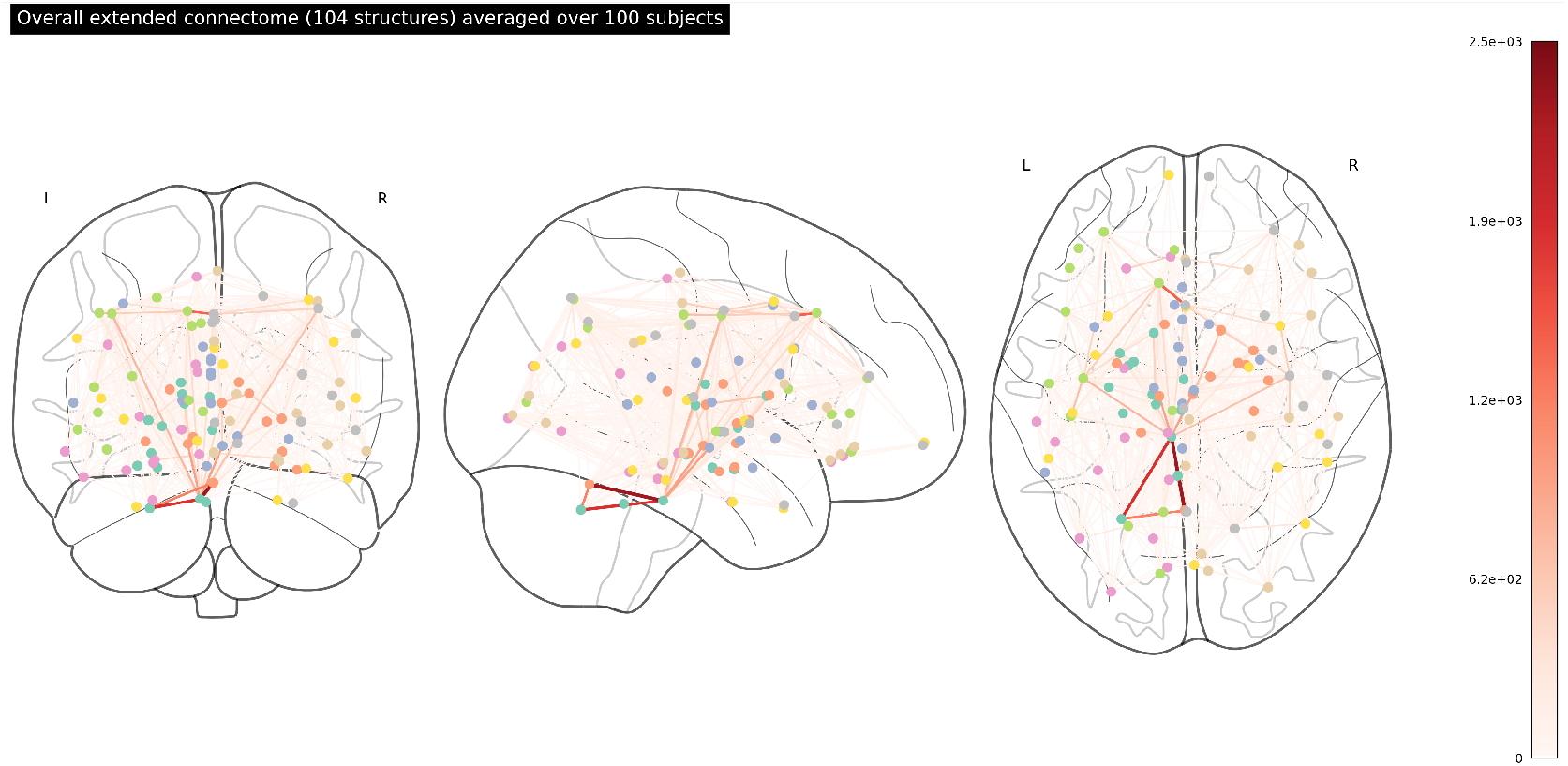
An overarching connectome featuring the 104 structures in the extended network. The strength of connections between structures (represented by the coloured nodes) is indicated by the colour, with darker red indicating more streamlines connecting two regions. Streamlines correspond to white matter fibre bundles. We note very strong edge-weighted connections between the left and right cerebellum cortices and the brainstem, as well as several strong connections in the midbrain and mid-cortex region. The full list of the 104 structures in this connectome can be found in **S1 Table**.

**Fig 2.**
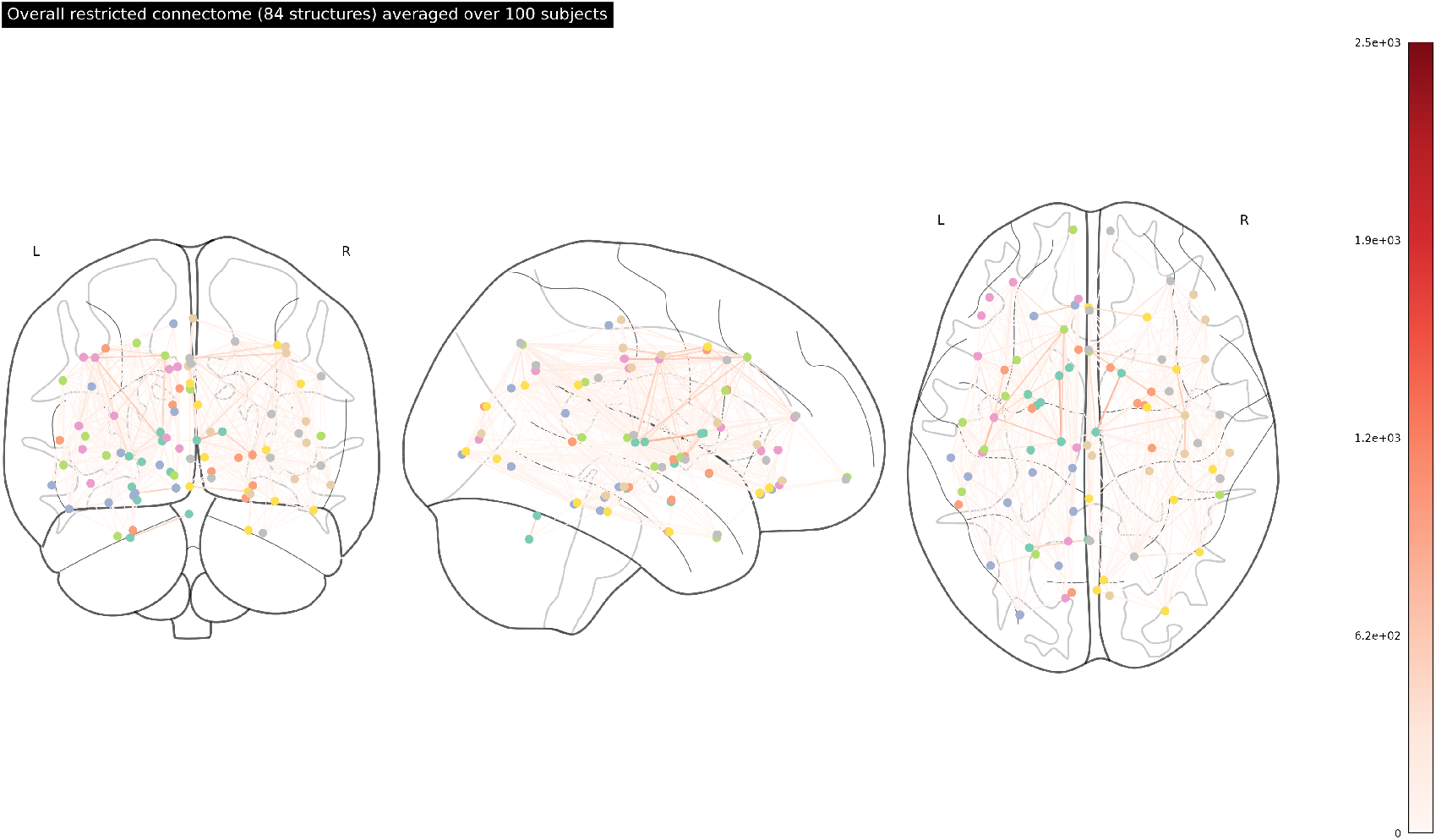
An overarching connectome featuring the 84 structures in the restricted network. With the absence of the brainstem, both sides of the cerebellum appear to be very weakly connected to the rest of the network. Additionally, we no longer see as many strong connections in the midbrain and mid-cortex region. The full list of the 84 structures in this connectome can be found in **S2 Table**.

**Fig 3.**
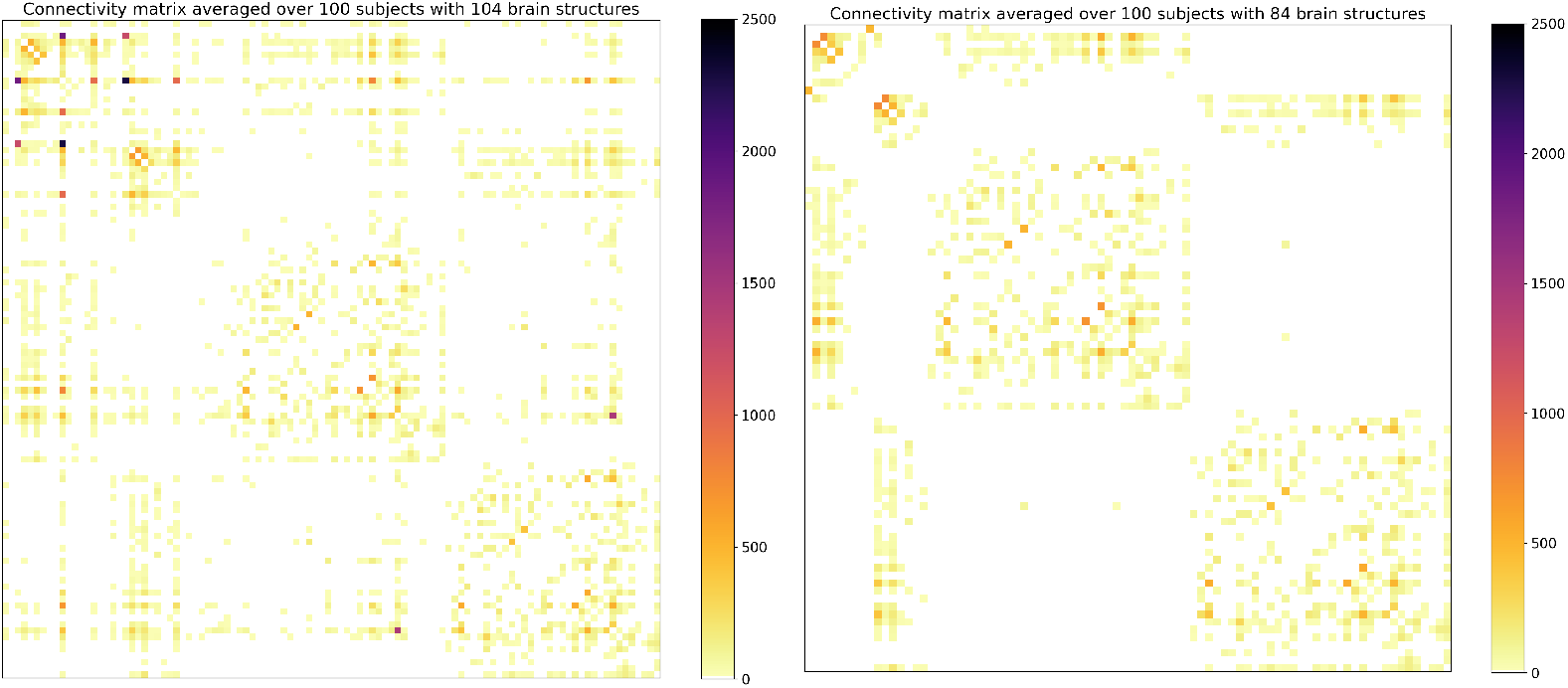
(a) (left) The structural connectivity matrix for the extended connectome (104 structures), averaged over 100 adult subjects; (b) (right) the structural connectivity matrix for the restricted connectome (84 structures). The full list of structures in the extended and restricted connectomes can be found in **S1 Table** and **S2 Table** respectively.

### Centrality

As explained above, we quantify the importance to the network of each node in three different ways; we compute the degree centrality, eigenvector centrality, and betweenness centrality. We present in Fig 4 the 10 highest ranked structures according to each of the three measures, both within the extended network (a) and the restricted one (b). Each centrality measure is normalized by dividing by its own maximum value, such that the highest ranked structure according to each measure always has a value of unity. In order to take into account the variable size of the structures, we performed the same calculations for normalized versions of each network. The extended and restricted networks were both normalized by dividing the strength of each connection by the sum of the volumes (lists of which are presented in **S1 Table** and **S2** Table) of its two nodes, as in earlier studies [45, 46]. The results for the normalized networks are presented in Fig 5.

**Fig 4.**
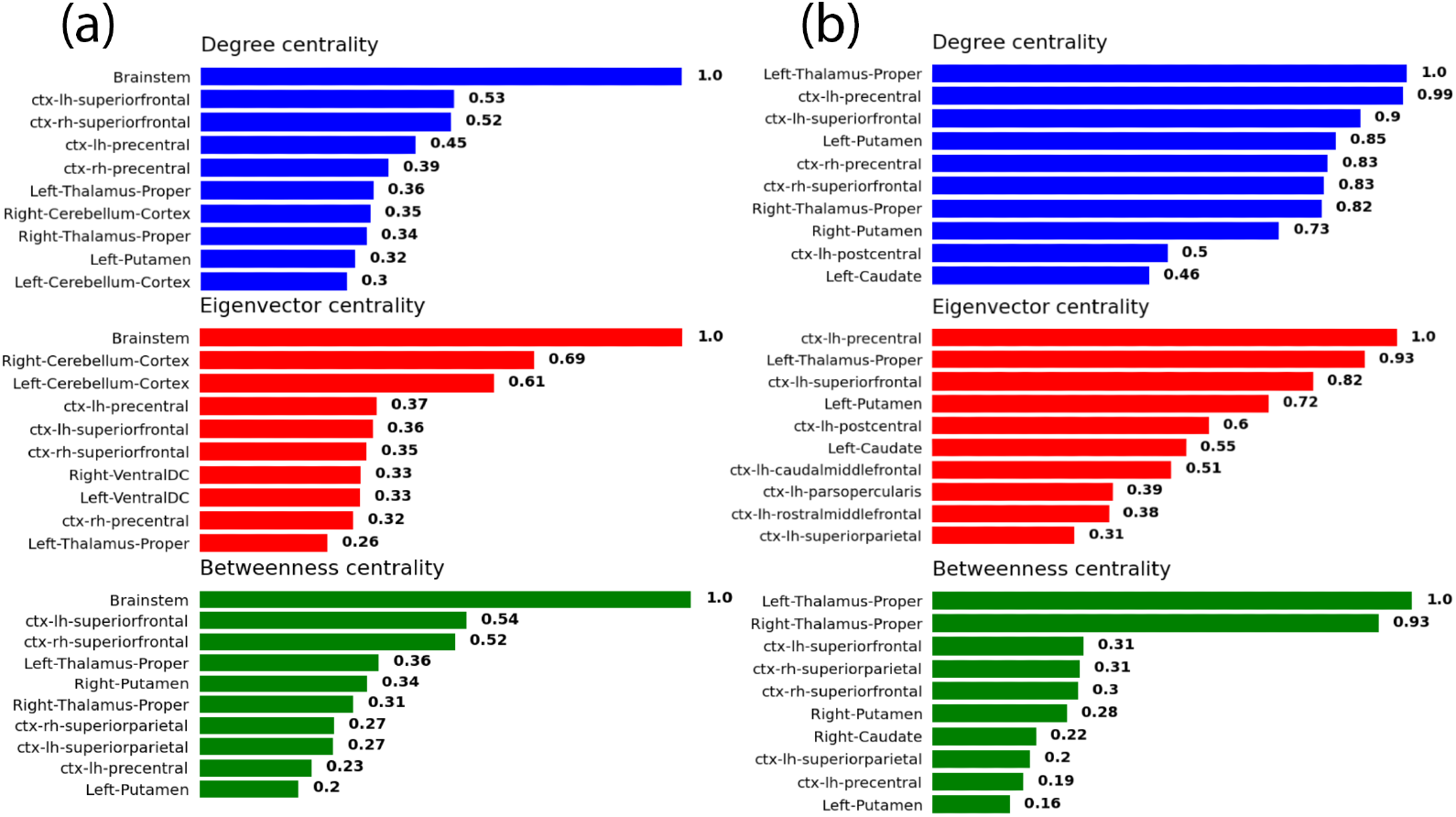
The top 10 central structures within the averaged connectome, for (a) the extended network and (b) the restricted network, according to three different measures: degree centrality (top), eigenvector centrality (middle), and betweenness centrality (bottom). Each of the three quantities is normalized with respect to its own maximum value.

**Fig 5.**
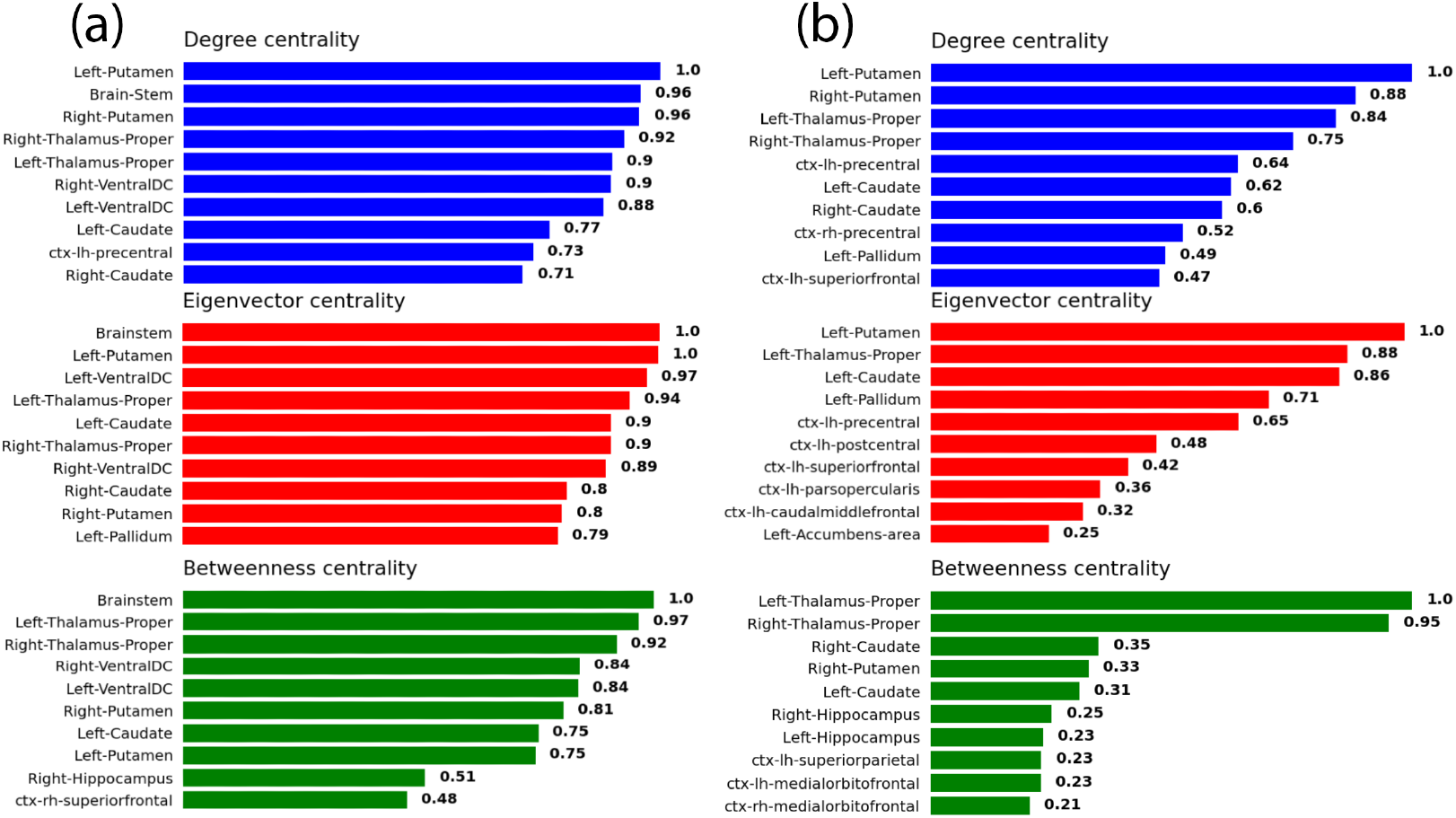
The top 10 central structures within the averaged *volume-normalized* connectome, for (a) the extended network and (b) the restricted network, according to three different measures: degree centrality (top), eigenvector centrality (middle), and betweenness centrality (bottom). Each of the three quantities is normalized with respect to its own maximum value.

We begin by discussing the case of the extended connectome in Fig 4a. In all the three centralities, the brainstem is leading by quite a substantial margin. This result clearly establishes the high centrality of the brainstem within the averaged connectome, at least from the perspective of the three measures considered here. Another structure which displays strong centrality (albeit nowhere near as strong as the brainstem) is the superior frontal cortex, as both the left and right hand-sides feature in the high ranks of all three measures.

The remaining structures featured in Fig 4a, while still displaying considerable centrality, pale in prominence compared to the brainstem, and even to the superior frontal cortex. Some structures are ranked highly according to two measures but not the third, such as the thalamus and the precentral cortex. Other high-ranking structures of lesser importance include the ventral diencephalon (DC), the putamen, the cerebellum, and the superior parietal cortex.

Fig 4b paints quite a different picture for the restricted connectome. A particularly striking feature, for instance, is the absence of a structure that strongly dominates all others, in the same way that the brainstem does in Fig 4a. In its stead, the most salient subcortical structure appears to be the thalamus, which, while ranking rather highly in general, only appears to eminently dominate in the case of the betweenness centrality. In the cases of degree and eigenvector centralities, there is a notable absence of a single structure that dwarves the rest.

Another noteworthy difference between Fig 4a and b is the order in which the structures appear. It is clear that, going from the extended connectome to the restricted one, the surviving structures do not retain the order of their centralities. The removal of those 20 structures has indeed caused some structures to rise in centrality, and others to fall. This is also the case for the corresponding normalized networks in Fig 5a and b. However, some structures such as the thalamus and the putamen retain their importance in both Figs 4 and 5, when going from the extended network (a) to the corresponding restricted one (b).

The cerebellum is another sub-cortical structure which, at first glance, may appear to possess a considerable amount of degree and eigenvector centrality within the extended connectome in Fig 4a. However, when the volume density of streamlines is taken into account, the situation is altered. Consider Fig 5a, which does not feature the cerebellum among any of the high ranking centrality measures. In fact, the degree, eigenvector, and betweenness centralities for the cerebellum in this case were computed to be 0.14, 0.18, and 0 respectively. This strongly suggests that the apparent high connectivity of the cerebellum implied by Fig 4a may largely be attributable to its sheer size. This is not true of the brainstem, on the other hand, which retains quite a high ranking in Fig 5a across all centrality measures, despite its substantial volume. The superiorfrontal cortex is another example of a structure that loses some prominence when the volume is taken into account, albeit to a much lesser extent than the cerebellum.

## Discussion

Our results unambiguously highlight the structural importance of the brainstem in the extended connectome. All measures of centrality we computed point to the brainstem being the most prominent structure in the averaged network, by a margin larger than the differences between the successive structures. It also became clear upon studying the volume-normalized network that the large size of the brainstem is not solely responsible for its dominance. By contrast, it appears that the high centrality of the cerebellum in the non-normalized network was an artifact of its extensive volume.

Furthermore, the connections of the cerebellum reach primarily to the brainstem, and to very few structures in the midbrain and beyond in the cortex. This becomes clear when comparing the ranking of the cerebellum between Fig 4a, with all 104 structures present, and 4b, with 20 structures (including the brainstem) absent. Upon exclusion of the brainstem, the cerebellum disappears completely from the list of top 10 structures for all three centrality measures. This, in combination with the volume observation, certainly detracts from the importance of the cerebellum within the structural network, which is at least consistent with the accepted idea that the cerebellum does not take any part in conscious processing [47].

As mentioned in the results section, going from the extended to the restricted connectome did not preserve the centrality order of the surviving structures. While perhaps unsurprising, this observation emphasizes the importance of incorporating as many brain regions as possible when drawing conclusions based on such graph-theoretical considerations. For example, the use of an exclusively cortical atlas, even if only cortical structures are of interest, may yield misleading results.

Obviously, the computed centrality measures are not identically defined, and thus can hardly be expected to agree on the ranking of every brain region. It was therefore unsurprising to find that some structures, such as the superior parietal cortex, ranked highly in some centrality measures but not others. However, one observation for which we do not find a straightforward explanation was the dominance of the left hemisphere in the eigenvector centrality rankings *only* in the case of the restricted connectome. While a possible explanation for this observation could be the fact that the majority of people are right-handed, this still would not explain why the dominance is only observed when several subcortical structures are removed. It is rather surprising that the exclusion of these 20 structures should give rise to such an extensive asymmetry.

With regards to theories of consciousness, the high ranking of the brainstem across all examined metrics suggests an important role for the brainstem in information integration. Recent studies suggest that mechanisms in the brainstem are crucial for the constitution of the conscious state [30]. At the very least, the brainstem is responsible for maintaining a most basic level of consciousness, as evidenced by various studies on children with hydranencephaly, who still appeared conscious despite missing most of their cortex [30]. Damage to the brainstem can induce a comatose state, which is the most radical form of disturbance in consciousness [48]. Given the evidence linking the brainstem to important aspects of consciousness, and our results suggesting the brainstem is one of the most, if not the most, highly connected anatomical region in the brain, we propose that the brainstem warrants more consideration within notable theories of consciousness. With some theorists suggesting that the upper brainstem may be especially important for maintaining the state of consciousness ([29], [28]), this also calls for more structural network analyses with finer parcellations of the brainstem.

Global workspace theory proposes a fleeting memory capacity that enables access between brain functions [21]. While it is a functional theory, it does draw on a structurally inspired framework [49], and it is thought that the thalamus and cortex work as an integrated system to create this global access network, because they are functionally integrated to a degree where they constitute a single functional system [50]. Our metrics suggest that the superior frontal and precentral cortices are strongly connected to the rest of the network, as are the right and left thalami, which suggests that a strong structural cortico-thalamic network may be present. Although research on brainstem functional connectivity is limited and the brainstem is typically not considered in GWT, there is new evidence to suggest that a brainstem-cortical interplay is critical for consciousness [51], which is supported by our results. We see a heavy influence of the brainstem in our structural network, which we argue warrants a deeper investigation into brainstem functional connectivity within the framework of GWT.

The postulates of IIT attempt to identify a physical substrate of consciousness (PSC), where the full NCCs correspond to neural structures that constitute the PSC, independent of their functional state [52]. IIT further defines consciousness as the capacity of a system to integrate information [53], which is quantified by *ϕ*_*max*_. To achieve a high *ϕ*_*max*_, a structure must be sufficiently functionally integrated [54]. However, there is evidence to suggest a correlation between structural and functional connectivity, and given that the PSC can split as a result of anatomical disconnections [52], there is reason to believe that structural connectivity does play a fundamental role in IIT. Patients with disorders of consciousness (DOC), where their anatomical connectivity is significantly altered, have been reported to have an associated reduction in functional connectivity [55]. Examining our results in the framework of IIT, under the assumption that structural and functional connectivity are related, we see a likelihood for the brainstem to give rise to a high *ϕ*_*max*_ given that it is the top-scoring structure across all our structural metrics. This provides further motivation to examine the brainstem from a functional perspective in the context of IIT as well.

The thalamacortical loop theory is another prominent functional theory of consciousness, where the thalamus acts as a hub which modulates communication with the cortex [56]. It proposes that consciousness is determined by synchronous gamma spikes in the thalamocortical system [56], meaning that the thalamocortical circuit dynamics contribute to certain NCCs [57]. Although it is a functional theory, overlapping patterns of structural and functional connectivity have been observed in the thalamic nuclei [58], suggesting a role for structural connectivity within the thalamocortical framework as well. In DOC patients for example, structural connections between the thalamic nuclei, the pallidum, and the frontal cortices have been shown to decrease [59]. Our results thus show that certain cortical structures as well as the thalamus are also very well connected, providing structural evidence for a thalamocortical network.

Finally, we make a connection between the results presented here and the newly emerging ideas involving quantum effects in the brain (see, for instance, [60]). Given that the delivery of anaesthetics specifically targeting the brainstem has been shown to be sufficient for the induction of loss of consciousness [61], and given the recent proposal that quantum effects may be important for xenon-induced anaesthesia [62], the prominent role enjoyed by the brainstem in our structural analysis may have potential implications for quantum-mechanical explanations of consciousness.

There are several limitations with our computational fibre tracking method. One of the largest limitations with fibre tractography using diffusion tensor imaging (DTI) is that the tensor model, which characterizes the orientation of white matter fibres, cannot differentiate complex configurations of crossing fibres in a single voxel [63]. Although the incorporation of constrained spherical deconvolution (CSD) in our computation improves this error, there are a few other advanced computational methods available, such as Q-ball imaging and diffusion spectrum imaging, and there is no general consensus on which method is best [63]. The CSD uses an empirically derived response function, which estimates the direction of fibre tracts and assumes that white matter fibres do not differ in their diffusion properties between tracts [64]. A common criticism of this approach, however, is that the response function may not be constant throughout the brain [64], which may alter the measured distribution of fibres within a voxel. Additionally, while there is no clear consensus on which parcellation method is best, the structures extracted from the HCP structural images used in this paper do not allow for a more detailed examination of the brainstem, as it has not been parcellated into smaller regions such as the upper nuclei, medulla, and pons.

The CSD approach is most notably used in MRtrix, a widely-used software pipeline for streamline tractography and connectome generation. It is formulated such that either a probabilistic or deterministic tracking algorithm can be used to track and generate streamlines. The `tckgen` function uses a probabilistic tracking algorithm by default, taking a fibre orientation distribution (FOD) image as input, where streamlines are considered more probable to follow a path where the FOD amplitudes along that path are also large. Similarly, our Python approach uses CSD to generate an FOD that guides the `ProbabilisticDirectionGetter` in the DIPY module, producing tracking direction seeds that are then used in the tracking. The difference between the methods, however, lies in their approaches to track termination. MRtrix terminates a track during the tracking process if no suitable FOD peak can be found whose amplitude is above a user-specified threshold [65]. In Python, we first used a Constant Solid Angle (CSA) Orientation Distribution Function (ODF) model, or the `CsaOdfModel` function, to estimate the orientation of tract segments at a point in the image. Then, the CSA/ODF model was passed into the `ThresholdStoppingCriterion` function, which determines the stopping criterion for the tracks based on the peaks of the ODF. Since the Desikan-Killiany atlas, which produces the restricted connectome, is available for use with MRtrix, we ran the software on one subject to compare the two methods. We noticed some quantitative differences, as some structures appeared to be higher or lower scoring in the MRtrix version than in our Python version, but qualitatively, the matrices were similar. Largely the same structures appeared in the top ten across all our centrality measures. An additional key difference between the two methods is that MRtrix allows for the manual setting of the number of streamlines desired, whereas in DIPY, the number is determined empirically based on a subject’s structural data. This renders it difficult to conduct a direct quantitative comparison between the two methods, since the convergence of the MRtrix results as a function of the number of streamlines may be more ambiguous.

Nevertheless, our qualitative comparison shows there can be differences in the strength of certain structures depending on the computational methods used. An extension of our work could involve computing the same centrality measures for an MRtrix-generated connectome if an atlas including both cortical structures and subcortical structures (including the brainstem) should become available, and to quantify the optimal number of streamlines in MRtrix.

Our computed centrality measures, while powerful tools, also come with some limitations. Degree centrality, for instance, straightforward as it is to compute, assumes that the importance of a node is determined entirely by the amount of connections it has to other nodes in the network. Hence, such a measure could in principle overestimate the importance of a node which is connected to a large number of irrelevant nodes. To rectify this shortcoming, eigenvector centrality gives precedence to those nodes that are connected to other nodes with high centrality, as opposed to considering all connections on equal grounds. One possible limitation of eigenvector centrality is that it says little of the difference between the degree of a node and those of its neighbours, which is an important consideration for the study of assortativity and modularity of networks [66].

These two measures are complemented by betweenness centrality, which identifies the most important nodes of a network as those which lie in the most strategic locations with respect to the shortest paths between pairs of nodes. While certainly a strong measure, it suffers from its own limitations, such as a computational cost that quickly becomes unfeasible as a function of network size. Although our moderately sized networks allowed us to largely evade this limitation in this work, it is bound to present a challenge for future studies considering significantly larger brain networks. Furthermore, betweenness centrality is implicitly based on the assumption that information propagates through the network serially along the most efficient routes. This has been argued to be a flawed assumption when considering brain networks, as it presupposes that individual components possess global information about the network topology [67], a condition that is unlikely to be satisfied in the context of biological systems [68].

The notion of network directionality constitutes another limitation within our approach. Graph-theoretical studies based on neuroimaging data typically represent connections within networks as undirected, owing to the inability of such non-invasive techniques to resolve the directionality of anatomical connections [17]. Our investigation presented here is no exception. However, it is believed that brain network connections are inherently directional in nature, and that the assumption of undirected connections introduces inaccuracies to the computation of topological properties, especially to the process of hub identification [69]. Hence, results obtained here may in principle be improved in future studies by constructing directed networks. Currently, some invasive techniques are capable of resolving axonal directionality of non-human connectomes [68].

A natural extension of this research would be to examine the role of the brainstem in more detail, ideally with a finer parcellation of the entire connectome. The brainstem could be parcellated into the midbrain, pons, medulla, and superior cerebellar peduncles [70]. This may alter the structural network even further. Another topic for future investigation is establishing connections with functional observations by means of implementing a dynamical model in the context of our network. A number of such models have been applied to brain networks to assess structure-function relationships, such as neural mass models [71, 72], models of Wilson-Cowan oscillators [46, 73], and spin models [31, 32, 33]. Studying critical phenomena and phase transitions within these systems allows for the possibility of making connections with the critical brain hypothesis [74, 75].

## Conclusion

We examined structural connectivity across 100 healthy adult subjects, using a specialized computational approach that includes the brainstem. We found that the brainstem scores the highest in all our computed centrality measures, followed by the superiorfrontal cortices and the right and left thalami. When the brainstem and several other subcortical structures are removed in the restricted connectome, we found that the most prominent structures revealed by the centrality analysis were quite different. This stresses the importance of incorporating all brain regions in structural network analyses, and suggests that the use of a non-comprehensive list of structures may give rise to misleading conclusions. Given evidence suggesting that structural connectivity plays a role in the modulation of conscious processing, our results suggest that subcortical structures, especially the brainstem, merit more consideration when examining theories of consciousness.

## Supporting information

**S1 Table.** A list of the 104 brain structures used in the extended connectivity matrix, with the corresponding number of streamlines attached to each structure and the average volume of the structure across 100 subjects.

| ID | Brain structure | Number of streamlines connected to the structure | Structure volume ($mm^3$) |
| --- | --- | --- | --- |
| 1 | Left-Lateral-Ventricle | 1190 | 6443.281 |
| 2 | Left-Inf-Lat-Vent | 45 | 235.092 |
| 3 | Left-Cerebellum-Cortex | 3939 | 57968.859 |
| 4 | Left-Thalamus-Proper | 4670 | 8619.509 |
| 5 | Left-Caudate | 2377 | 3851.667 |
| 6 | Left-Putamen | 4204 | 5540.322 |
| 7 | Left-Pallidum | 1552 | 1359.246 |
| 8 | 3rd-Ventricle | 348 | 782.034 |
| 9 | 4th-Ventricle | 434 | 1825.385 |
| 10 | Brain-Stem | 13150 | 22167.976 |
| 11 | Left-Hippocampus | 852 | 4564.249 |
| 12 | Left-Amygdala | 190 | 1593.61 |
| 13 | CSF | 362 | 1063.154 |
| 14 | Left-Accumbens-area | 384 | 573.736 |
| 15 | Left-VentralDC | 3784 | 4292.068 |
| 16 | Left-vessel | 102 | 59.189 |
| 17 | Left-choroid-plexus | 554 | 1125.245 |
| 18 | Right-Lateral-Ventricle | 978 | 5884.33 |
| 19 | Right-Inf-Lat-Vent | 38 | 232.622 |
| 20 | Right-Cerebellum-Cortex | 4659 | 59587.369 |
| 21 | Right-Thalamus-Proper | 4573 | 7521.443 |
| 22 | Right-Caudate | 2378 | 3952.693 |
| 23 | Right-Putamen | 4037 | 5609.062 |
| 24 | Right-Pallidum | 658 | 1493.601 |
| 25 | Right-Hippocampus | 985 | 4627.455 |
| 26 | Right-Amygdala | 226 | 1655.291 |
| 27 | Right-Accumbens-area | 435 | 600.805 |
| 28 | Right-VentralDC | 3927 | 4299.531 |
| 29 | Right-vessel | 148 | 71.884 |
| 30 | Right-choroid-plexus | 500 | 1235.485 |
| 31 | Optic-Chiasm | 264 | 231.912 |
| 32 | CC_Posterior | 231 | 960.037 |
| 33 | CC_Mid_Posterior | 316 | 478.638 |
| 34 | CC_Central | 436 | 514.796 |
| 35 | CC_Mid_Anterior | 417 | 515.937 |
| 36 | CC_Anterior | 342 | 900.829 |
| 37 | ctx-lh-bankssts | 192 | 2862.336 |
| 38 | ctx-lh-caudalanteriorcingulate | 557 | 3022.485 |
| 39 | ctx-lh-caudalmiddlefrontal | 1967 | 7065.912 |
| 40 | ctx-lh-cuneus | 467 | 2200.646 |
| 41 | ctx-lh-entorhinal | 268 | 865.957 |
| 42 | ctx-lh-fusiform | 762 | 7097.376 |
| 43 | ctx-lh-inferiorparietal | 1090 | 10072.529 |
| 44 | ctx-lh-inferiortemporal | 612 | 6603.035 |
| 45 | ctx-lh-isthmuscingulate | 927 | 3957.884 |
| 46 | ctx-lh-lateraloccipital | 816 | 8963.47 |
| 47 | ctx-lh-lateralorbitofrontal | 1029 | 6725.032 |
| 48 | ctx-lh-lingual | 773 | 5273.808 |
| 49 | ctx-lh-medialorbitofrontal | 1528 | 3930.844 |
| 50 | ctx-lh-middletemporal | 603 | 5530.654 |
| 51 | ctx-lh-parahippocampal | 299 | 1668.477 |
| 52 | ctx-lh-paracentral | 966 | 3732.204 |
| 53 | ctx-lh-parsopercularis | 1629 | 3869.169 |
| 54 | ctx-lh-parsorbitalis | 183 | 867.128 |
| 55 | ctx-lh-parstriangularis | 642 | 3038.14 |
| 56 | ctx-lh-pericalcarine | 482 | 3075.872 |
| 57 | ctx-lh-postcentral | 2426 | 7322.379 |
| 58 | ctx-lh-posteriorcingulate | 878 | 4473.498 |
| 59 | ctx-lh-precentral | 5905 | 13190.15 |
| 60 | ctx-lh-precuneus | 1658 | 9317.325 |
| 61 | ctx-lh-rostralanteriorcingulate | 500 | 2617.448 |
| 62 | ctx-lh-rostralmiddlefrontal | 1974 | 12890.373 |
| 63 | ctx-lh-superiorfrontal | 6817 | 18488.612 |
| 64 | ctx-lh-superiorparietal | 2264 | 12374.398 |
| 65 | ctx-lh-superiortemporal | 946 | 7718.647 |
| 66 | ctx-lh-supramarginal | 1003 | 8872.386 |
| 67 | ctx-lh-frontalpole | 134 | 190.543 |
| 68 | ctx-lh-temporalpole | 200 | 661.269 |
| 69 | ctx-lh-transversetemporal | 335 | 807.284 |
| 70 | ctx-lh-insula | 1159 | 8607.619 |
| 71 | ctx-rh-bankssts | 179 | 2950.619 |
| 72 | ctx-rh-caudalanteriorcingulate | 673 | 3172.229 |
| 73 | ctx-rh-caudalmiddlefrontal | 2115 | 6180.176 |
| 74 | ctx-rh-cuneus | 503 | 2191.445 |
| 75 | ctx-rh-entorhinal | 360 | 661.167 |
| 76 | ctx-rh-fusiform | 794 | 6957.369 |
| 77 | ctx-rh-inferiorparietal | 1265 | 12013.279 |
| 78 | ctx-rh-inferiortemporal | 629 | 6112.701 |
| 79 | ctx-rh-isthmuscingulate | 666 | 3573.922 |
| 80 | ctx-rh-lateraloccipital | 705 | 8876.928 |
| 81 | ctx-rh-lateralorbitofrontal | 1246 | 6661.849 |
| 82 | ctx-rh-lingual | 723 | 5407.399 |
| 83 | ctx-rh-medialorbitofrontal | 1317 | 3406.247 |
| 84 | ctx-rh-middletemporal | 807 | 6306.081 |
| 85 | ctx-rh-parahippocampal | 350 | 1680.065 |
| 86 | ctx-rh-paracentral | 1267 | 4657.647 |
| 87 | ctx-rh-parsopercularis | 1071 | 3563.056 |
| 88 | ctx-rh-parsorbitalis | 274 | 1191.086 |
| 89 | ctx-rh-parstriangularis | 651 | 3366.89 |
| 90 | ctx-rh-pericalcarine | 512 | 3168.029 |
| 91 | ctx-rh-postcentral | 2041 | 7181.13 |
| 92 | ctx-rh-posteriorcingulate | 883 | 4517.583 |
| 93 | ctx-rh-precentral | 5109 | 13710.037 |
| 94 | ctx-rh-precuneus | 1627 | 10019.167 |
| 95 | ctx-rh-rostralanteriorcingulate | 376 | 2153.274 |
| 96 | ctx-rh-rostralmiddlefrontal | 1827 | 13480.068 |
| 97 | ctx-rh-superiorfrontal | 6864 | 18600.449 |
| 98 | ctx-rh-superiorparietal | 2516 | 11960.449 |
| 99 | ctx-rh-superiortemporal | 1019 | 6898.772 |
| 100 | ctx-rh-supramarginal | 953 | 8860.182 |
| 101 | ctx-rh-frontalpole | 234 | 285.97 |
| 102 | ctx-rh-temporalpole | 279 | 613.389 |
| 103 | ctx-rh-transversetemporal | 258 | 576.932 |
| 104 | ctx-rh-insula | 1510 | 8912.524 |

**S2 Table.** A list of the 84 brain structures used in the restricted connectivity matrix, with the corresponding number of streamlines attached to each structure.

| Brain structure | Number of streamlines connected to the structure |
| --- | --- |
| Left-Cerebellum-Cortex | 529 |
| Left-Thalamus-Proper | 3818 |
| Left-Caudate | 1740 |
| Left-Putamen | 3239 |
| Left-Pallidum | 999 |
| Left-Hippocampus | 383 |
| Left-Amygdala | 74 |
| Left-Accumbens-area | 277 |
| Right-Cerebellum-Cortex | 490 |
| Right-Thalamus-Proper | 3129 |
| Right-Caudate | 1712 |
| Right-Putamen | 2791 |
| Right-Pallidum | 526 |
| Right-Hippocampus | 502 |
| Right-Amygdala | 88 |
| Right-Accumbens-area | 313 |
| ctx-lh-bankssts | 202 |
| ctx-lh-caudalanteriorcingulate | 532 |
| ctx-lh-caudalmiddlefrontal | 1649 |
| ctx-lh-cuneus | 390 |
| ctx-lh-entorhinal | 261 |
| ctx-lh-fusiform | 732 |
| ctx-lh-inferiorparietal | 994 |
| ctx-lh-inferiortemporal | 633 |
| ctx-lh-isthmuscingulate | 758 |
| ctx-lh-lateraloccipital | 639 |
| ctx-lh-lateralorbitofrontal | 898 |
| ctx-lh-lingual | 578 |
| ctx-lh-medialorbitofrontal | 1068 |
| ctx-lh-middletemporal | 598 |
| ctx-lh-parahippocampal | 256 |
| ctx-lh-paracentral | 425 |
| ctx-lh-parsopercularis | 1488 |
| ctx-lh-parsorbitalis | 162 |
| ctx-lh-parstriangularis | 564 |
| ctx-lh-pericalcarine | 399 |
| ctx-lh-postcentral | 1898 |
| ctx-lh-posteriorcingulate | 762 |
| ctx-lh-precentral | 3784 |
| ctx-lh-precuneus | 1089 |
| ctx-lh-rostralanteriorcingulate | 409 |
| ctx-lh-rostralmiddlefrontal | 1456 |
| ctx-lh-superiorfrontal | 3451 |
| ctx-lh-superiorparietal | 1612 |
| ctx-lh-superiortemporal | 878 |
| ctx-lh-supramarginal | 960 |
| ctx-lh-frontalpole | 89 |
| ctx-lh-temporalpole | 192 |
| ctx-lh-transversetemporal | 333 |
| ctx-lh-insula | 1110 |
| ctx-rh-bankssts | 179 |
| ctx-rh-caudalanteriorcingulate | 593 |
| ctx-rh-caudalmiddlefrontal | 1661 |
| ctx-rh-cuneus | 409 |
| ctx-rh-entorhinal | 315 |
| ctx-rh-fusiform | 748 |
| ctx-rh-inferiorparietal | 1156 |
| ctx-rh-inferiortemporal | 605 |
| ctx-rh-isthmuscingulate | 553 |
| ctx-rh-lateraloccipital | 570 |
| ctx-rh-lateralorbitofrontal | 956 |
| ctx-rh-lingual | 580 |
| ctx-rh-medialorbitofrontal | 849 |
| ctx-rh-middletemporal | 711 |
| ctx-rh-parahippocampal | 306 |
| ctx-rh-paracentral | 671 |
| ctx-rh-parsopercularis | 1010 |
| ctx-rh-parsorbitalis | 236 |
| ctx-rh-parstriangularis | 574 |
| ctx-rh-pericalcarine | 431 |
| ctx-rh-postcentral | 1559 |
| ctx-rh-posteriorcingulate | 780 |
| ctx-rh-precentral | 3178 |
| ctx-rh-precuneus | 1040 |
| ctx-rh-rostralanteriorcingulate | 290 |
| ctx-rh-rostralmiddlefrontal | 1306 |
| ctx-rh-superiorfrontal | 3154 |
| ctx-rh-superiorparietal | 1738 |
| ctx-rh-superiortemporal | 868 |
| ctx-rh-supramarginal | 878 |
| ctx-rh-frontalpole | 139 |
| ctx-rh-temporalpole | 238 |
| ctx-rh-transversetemporal | 256 |
| ctx-rh-insula | 1322 |

**S3 Table.** The list of 20 structures removed in the restricted connectivity matrix. This matrix corresponds with the standard MRtrix parcellation scheme, widely used in studies of brain networks. The list of cortical structures remained the same.

| Removed Structures |
| --- |
| Left-Lateral-Ventricle |
| Left-Inf-Lat-Vent |
| 3rd-Ventricle |
| 4th-Ventricle |
| Brain-Stem |
| CSF |
| Left-VentralDC |
| Left-vessel |
| Left-choroid-plexus |
| Right-Lateral-Ventricle |
| Right-Inf-Lat-Vent |
| Right-VentralDC |
| Right-vessel |
| Right-choroid-plexus |
| Optic-Chiasm |
| CC_Posterior |
| CC_Mid_Posterior |
| CC_Central |
| CC_Mid_Anterior |
| CC_Anterior |

## Acknowledgments

This work was supported by the Natural Sciences and Engineering Research Council (NSERC) of Canada. We would also like to thank Emma Towlson and Wilten Nicola for their valuable input, guidance, and discussion, as well as Joern Davidsen, Davor Curic, and Omid Khajehdehi from the Complexity Science Group (CSG). Additionally, we thank the HCP for providing access to their data.

## Author Contributions

**Conceptualization:** Christoph Simon, Salma Salhi, Youssef Kora, Hadi Zadeh Haghighi

**Data curation:** Salma Salhi

**Formal analysis:** Youssef Kora

**Funding acquisition:** Christoph Simon

**Investigation:** Salma Salhi, Youssef Kora

**Methodology:** Salma Salhi, Youssef Kora, Gisu Ham

**Project Administration:** Christoph Simon, Youssef Kora

**Software:** Gisu Ham, Salma Salhi, Youssef Kora

**Supervision:** Christoph Simon

**Validation:** Salma Salhi, Youssef Kora, Christoph Simon

**Visualization:** Salma Salhi, Youssef Kora

**Writing - original draft preparation:** Salma Salhi, Youssef Kora

**Writing - review and editing:** Salma Salhi, Youssef Kora, Christoph Simon, Hadi Zadeh Haghighi

## Data Availability Statement

The data used in this project was provided by the Human Connectome Project (HCP; Principal Investigators: Bruce Rosen, M.D., Ph.D., Arthur W. Toga, Ph.D., Van J. Weeden, MD). HCP funding was provided by the National Institute of Dental and Craniofacial Research (NIDCR), the National Institute of Mental Health (NIMH), and the National Institute of Neurological Disorders and Stroke (NINDS). HCP data are disseminated by the Laboratory of Neuro Imaging at the University of Southern California. Structural and diffusion MRI images from the HCP, as well as lists of extracted structures, bvals, and bvecs, were all used to process the data in our Python program. 100 subjects were used, all of which are part of the “WU-Minn HCP Data - 1200 Subjects” dataset. A complete list of subject names is available upon request. All scripts used to generate the connectomes and associated plots are available on our Github repository.

